# Hydrodynamics of gliding penguin flipper suggests the adjustment of sweepback with swimming speeds

**DOI:** 10.1101/2021.05.24.445327

**Authors:** Masateru Maeda, Natsuki Harada, Hiroto Tanaka

## Abstract

Hydrodynamic performance of a gliding penguin flipper (wing) considering the backward sweep was estimated with computational fluid dynamics (CFD) simulation. A flipper of a gentoo penguin (*Pygoscelis papua*) was 3D scanned, smoothed, and a numerical fluid mesh was generated. For accurate yet resource-saving computation, an embedded large-eddy simulation (ELES) methods was employed, where the flow near the flipper was solved with large-eddy simulation (LES) and flow far away from the flipper was solved with Reynolds-averaged Navier-Stokes (RANS). The relative flow speed was fixed at 2 m s^−1^, close to the typical foraging speed for the penguin species. The sweep angle was set to be 0°, 30°, and 60°, while the angle of attack was varied between −40° and 40°, both are within the realistic ranges in the wing kinematics measurement of penguins in an aquarium. It was revealed that a higher sweep angle reduces the lift slope, but the lift coefficient is unchanged at a high angle of attack. Drag coefficient was reduced across the angles of attack with increasing the sweep angles. The drag polars suggest the sweep angle may be adjusted with the change in swimming speed and anhedral (negative dihedral) angle to minimise drag while maintaining the vertical force balance to counteract the positive buoyancy. This will effectively expand the swimming envelope of the gliding penguin, similar to a flying counterpart such as swift.

## Introduction

Penguins are highly specialized swimmers among birds that use wings to propel themselves in the water. The wings, also called flippers, are quite different from those of the flying birds, where the overall wing is thicker and stiffer, and the feathers are filament-like (Louw, 1992). Understanding the hydrodynamic mechanism of penguin swimming is of fundamental importance for many aspects of penguin as they entirely rely on the flippers for any aquatic locomotion from everyday foraging to long-distance migration.

In the past couple of decades, the advancement in bio-logging has enabled us to see the swimming performance of penguins (e.g., Sato et al., 2002). The swimming mechanisms of penguins are, however, largely unexplored. There are a handful of papers on the measurements of penguin swimming and flipper performance in the wind tunnel (Clark and Bemis, 1979; Hui, 1988; Bannasch, 1995) but they were not combined with the actual penguins’ wing attitude in gliding.

Here we report the gliding performance of a gentoo penguin flipper with detailed three-dimensional wing morphology and realistic wing posture from the measurement of swimming. The lift and drag forces at various angles of attack and sweep angles were computed with computational fluid dynamics (CFD) simulation. The maximum lift-to-drag ratio was achieved when the flipper has a moderate sweep and angle of attack slightly before stall. Interestingly, when considering the trimmed condition for the forces acting on the penguin body, the large sweepback angle of the flipper is beneficial by minimising the drag while generating sufficient downward vertical force.

## Methods

Wing attitudes in glides were measured via underwater filming of penguin swimming in an aquarium. A 3D wing geometrical model was generated based on the scanned data and the computational fluid dynamics (CFD) simulation was performed, which was validated with a water tunnel test.

### Wing angles measurement in the aquarium

The wing postures of the swimming penguins were measured in an aquarium using underwater cameras (GoPro HERO6 Black and GoPro HERO7 Black, GoPro Inc, USA) and analysing the videos via direct linear transformation (DLT) with a commercial software DIPP-motion V/3D (Ditect Co., Ltd., Japan). From the dataset, four parameters were extracted: swimming speed measured at the body centre; flapping angle (dihedral angle), sweep angle, and angle of attack for each wing. The details of the measurement and analysis are described elsewhere (Harada et al., *under review*). In the current study, our definitions of the wing angles are almost the same as Harada et al., in that they are based on the coordinate transformation between the wing coordinates and the shoulder-fixed coordinates (which are simply the body coordinates translated to each shoulder), but the characteristic points to calculate the angles are slightly different. Flap angle *ϕ* and sweep angle *Λ* were defined using the wingtip point (as opposed to the leading-edge marker point in Harada et al.). On the other hand, the angle of attack was defined using the leading edge point at the 50% wing length position, exactly the same definition as the *α*outer in our previous paper (Harada et al., *under review*). From the all swimming dataset, we selected only the horizontal and straight gliding bouts (sequences). The “straight (no turning)” swimming was defined as the segment of swimming where “the yaw rate greater than ±30 deg s^−1^ does not persist for more than 0.3 s” and the “horizontal (level)” swimming was defined as the segment of swimming where “the vertical speed at the body centre greater than ±0.3 m s^−1^ does not persist for more than 0.3 s.”

### Computational fluid dynamics simulation

#### Wing 3D scanning and modelling

To obtain the three-dimensional shape, a wing (flipper) of a gentoo penguin *Pygoscelis papua* was scanned at the Nagasaki Penguin Aquarium (Nagasaki, Japan) on 29 September 2016 using a 3D scanner Artec Space Spider (Artec 3D, Luxembourg) with the 3D resolution of 0.1 mm. The penguin was standing still on the ground and the right wing was held horizontally by human hands while scanning; therefore, the shape may deviate to some extent from the actual shape during swimming. A few circular colour stickers were attached on the surface to help the registration (matching the 3D data across different scanning sequences) in the later processing. In addition, a weak plastic pinch was attached to the leading edge of the flipper, which protrudes from the thin wing and prevents the scanner from losing a thin object when transitioning between dorsal and ventral sides. The raw point cloud was processed with Artec Studio 12 Professional application, where the pinch was removed, registration was applied, and a triangular mesh was generated with “sharp fusion” command. The mesh (figure 1A, orange object) had several problems, e.g., lacking some wing surfaces and had rough surface due to noise, feathers, or stickers. A cleaner surface is necessary for the numerical mesh generation for CFD. Therefore, a 3D CAD Rhinoceros 5 (Robert McNeel & Associates, USA) was used to generate a smooth NURBS surface, where a planform view photo of the flipper was used for supplementing the outline for the proximal portion of the wing. The NetworkSrf command was used with four spanwise curves (figure 1A, blue lines) and 23 cross-sectional curves (figure 1A, grey lines). The surface (figure 1B) was exported in the IGES format.

**Figure 1.**
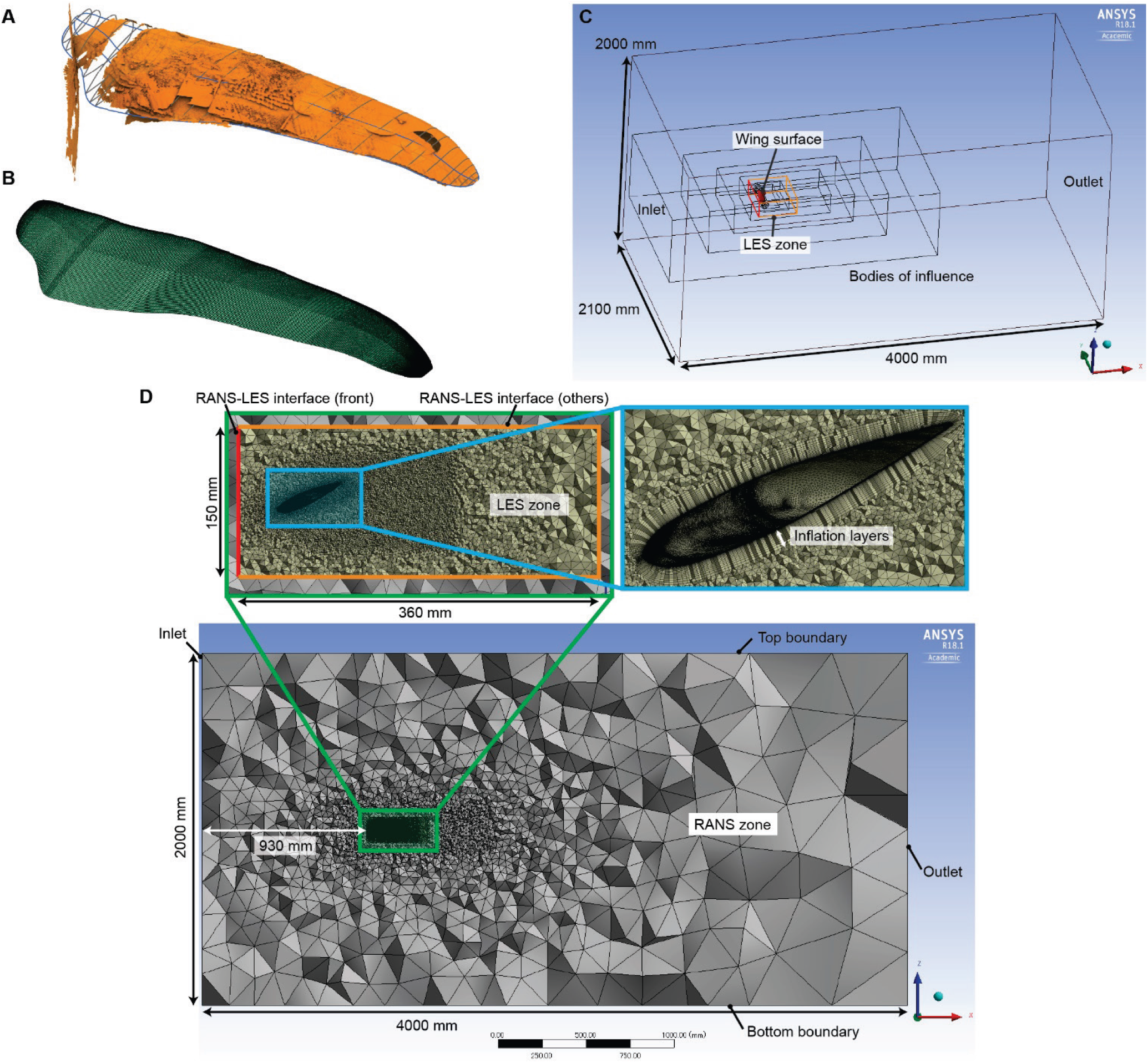
The morphological model of a penguin flipper, computational domain, and numerical mesh. The cross sections were extracted from the 3D scanned mesh (A). A smooth surface model was created using CAD software (B). Note a mirrored wing (left wing) was used for the flow computations. To show the mesh, a computational case of sweep angle *Λ* = 0°, α = −20° was chosen as an example. The screenshots are from mesh generation software ANSYS Meshing 18.1. The computational domain with dimensions (C). The red object in the middle is the wing surface. The outermost cuboid is the physical domain (4000 mm × 2100 mm × 2000 mm in X × Y × Z directions). A cuboid made of red and orange lines is the LES zone. Other objects inside of the physical domain are the virtual bodies for mesh size control (Bodies of Influence, BoI). Mesh at a cutting plane in the XZ-plane (D). The bottom panel shows the entire domain, where most of the volume is the RANS zone. The top left panel in (D) shows the enlarged view around the LES zone. The RANS-LES interfaces are coloured red (front) and orange (others). The top right panel in (D) shows the enlarged view around the wing model, showing the inflation layers (prism layers) to resolve the boundary layers. The discontinuous changes in mesh densities (except for the inflation layers) were achieved with the Bodies of Influence (4 for RANS zone, 6 for LES zone).

#### Numerical simulation setup

A flow domain for computational fluid dynamics was prepared using ANSYS DesignModeler (ANSYS, Inc., USA) in the Workbench 18.1 (figure 1C). For simplicity, the gliding condition around a single wing (flipper) was considered. The flipper surface generated in Rhinoceros was imported into DesignModeler. The small holes at the wing root and wingtip were closed, and the inner cavity was filled. The wing model was mirrored in the long-axis direction, i.e. effectively make it the left wing for aiming the better visualization. The wing attitude (sweep and feathering) was set for each case.

We chose a method called embedded large-eddy simulation (ELES) to minimize the computational resources while keeping the benefit of large-eddy simulation (LES). In short, a flow domain is divided into two zones: the outer zone solves Reynolds-averaged Navier-Stokes (RANS) while the inner zone (including the object surface) solves the LES. The flow variables at the interface are automatically interpolated. Therefore, for the outer zone, a large rectilinear box with the dimension of 4000 × 2100 × 2000 mm (length × width × height) was prepared. Another smaller box with 360 × 400 × 150 mm (length × width × height) was prepared near the wing for the LES zone. The inner box was subtracted from the outer (larger) box with the Boolean Subtract operation. Then, the wing was subtracted from the inner box with Boolean Subtract operation.

The positive x-axis is from the inlet face to the outlet face (from front to back). The positive y-axis is from left to right. The positive z-axis is from bottom to top. The wing long axis is defined as the line connecting the wingbase and the wingtip. The sweep angle *Λ* was defined to be 0 when the wing long axis is parallel to the y-axis. The wing has a moderate twist (i.e., each wing cross section has a slightly different angle of incidence), so we defined the angle of attack *α* = 0 when the lift is zero (not based on the morphology). The amount of shift was determined by several preliminary simulations. This procedure was repeated for each sweep angle; therefore, when the angle of attack *α* = 10° in a certain sweep angle and if you change the wing sweep, you won’t generally get the *α* = 10° at that swept posture.

#### Mesh generation

ANSYS Meshing in the Workbench 18.1 was used for numerical mesh generation (figure 1figure 1D). The mesh density is non-uniform so that the close to the wing the smaller the mesh size. The first layer thickness of the inflation layers (prism mesh for boundary layer), which is adjacent to the wing surface, was determined through several preliminary test computations to ensure *y*^+^ ≤ 1 for most of the wing surface, where *y*^+^ is defined as *y*^+^ ≔ (*ρu*_τ_*y*_p_)/*μ*, where *u*_*τ*_ ≔ *τ*_*w*_/*ρ* is the friction velocity, *y*_p_ is the distance from the boundary cell to the wall, *ρ* is the density of water, and *μ* is the dynamic viscosity of water (ANSYS Fluent Users Manual, version 18.1). The growth ratio and the number of layers for the inflation layer were also determined through test computations to ensure that the boundary layer (assumed to be twice the distance from the surface to the maximum turbulence viscosity ratio) is reasonably confined within those prism meshes. This is achieved by the technique called Body of Influence (BoI) in ANSYS Meshing. For this, 10 virtual bodies were placed around the wing model (4 for the RANS zone and 6 for the LES zone). Different mesh size was assigned for each virtual body, and this mesh size was kept nearly uniform within each virtual body. The size for the innermost virtual body was 1.1 mm, making the transition from the outermost layer of the inflation layers (thickness of 0.5 mm) relatively smooth. The maximum skewness for the mesh was confirmed to be less than 0.95 for each computational case. The total number of elements slightly vary between cases, but typically approximately 13 million.

#### Flow computation

ANSYS Fluent 18.1 was used for flow computation. The embedded large-eddy simulation (ELES) method was employed. The parameters and boundary conditions are summarized in Table 1. The uniform relative flow speed *U*_∞_ for the baseline simulation case was set at 2.0 m s^−1^ as this is close to the typical foraging swimming speed for several penguin species including the gentoo penguin (Sato et al., 2010). Note though, in contrast, penguins in aquarium seem to swim at a lower speed at around 1 m s^−1^ (e.g., Clark and Bemis, 1979; Harada et al., *under review*), although they do occasionally show rapid, burst swimming bouts. For each case, a steady RANS computation (without LES) was first performed until the flow was reasonably stabilised in terms of lift and drag. This usually took 1000 to 3000 iteration steps. When the flow is stabilised, a transient (unsteady) computation with the ELES method was initiated, where the final flow field in the RANS computation was used as the initial condition.

**Table 1.** Basic parameters. Summary of the wing and computation parameters. The density and viscosity of water are directly from the “water-liquid” material from the Fluent Database.

|  |  |
| --- | --- |
| Wing length, $r$ (mm) | 253 |
| Mean chord length, $c_m$ (mm) | 54.9 ($\Lambda = 0^\circ$), 63.4 ($\Lambda = 30^\circ$), 110 ($\Lambda = 60^\circ$) |
| Wing planform area, $S_w$ (mm<sup>2</sup>) | $1.39 \times 10^5$ |
| Density of water, $\rho$ (kg m<sup>-3</sup>) | 998.2 |
| Kinematic viscosity of water, $\nu$ (m<sup>2</sup> s<sup>-1</sup>) | $1.0048 \times 10^{-6}$ |
| Uniform flow speed, $U_\infty$ (m s<sup>-1</sup>) | 2.0 (baseline speed), 1.0 (lower speed), 4.0 (higher speed) |
| Reynolds number, Re | at baseline speed: $1.09 \times 10^5$ ($\Lambda = 0^\circ$), $1.26 \times 10^5$ ($\Lambda = 30^\circ$), $2.19 \times 10^5$ ($\Lambda = 60^\circ$)<br>at lower speed: $5.46 \times 10^4$ ($\Lambda = 0^\circ$), $1.09 \times 10^5$ ($\Lambda = 60^\circ$)<br>at higher speed: $2.19 \times 10^5$ ($\Lambda = 0^\circ$) |

There is a discontinuity of the mesh at the interface of virtual surfaces between the RANS zone (outer fluid domain) and the LES zone (inner fluid domain), where the interpolation of the flow variables between the inner and outer cells occurs. In addition, a velocity fluctuation is assigned to one of the interface surfaces, in front of the wing (figure 1C,D, red lines). We used the ‘Vortex Method’ for the velocity fluctuation and the number of vortices was calculated to be approximately 1/4 of the number of faces in the LES zone interface surface, as per the recommendation by the software developer (ANSYS, Inc.). The exact number of vortices differed slightly across cases as we generated mesh for each case (each wing angle), but it was approximately 500–600. The time step (or time increment, Δ*t*) for the transient computation was set as 0.0001 s for the baseline flow speed (*U*_∞_ = 2.0 m s^−1^) through a few preliminary computations to ensure the Courant number is overall ≤ 1.

#### Computation cases

For the baseline relative flow (swimming) speed (*U*_∞_ = 2.0 m s^−1^), the computations were carried out for sweep angles *Λ* = 0°, 30°, and 60° across the range of angles of attack −40° ≤ α ≤ 40°. In addition, a lower flow speed at *U*_∞_ = 1.0 m s^−1^ with two sweep angles (0°, and 60°) and several angles of attack (−40° ≤ α ≤ 40°, but a smaller number of sampling points) were computed. The inclusion of the negative angles of attack is particularly important for penguin flipper, as the generation of downward lift would be needed to counteract the positive buoyancy (the upward force vector resulting from the sum of weight and buoyancy forces) that is normally expected for penguins, with the deep dive as a potential exception. Furthermore, for the sweep angle *Λ* = 0°, a higher swimming speed (*U*_∞_ = 4.0 m s^−1^) was also computed.

#### Forces

The results are summarized by the force coefficients, drag polar, and lift-to-drag ratio. The force coefficients are obtained in the following equations:

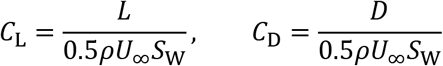

where *C*_L_ and *C*_D_ are lift coefficient and drag coefficient, respectively; *L* and *D* are lift and drag, respectively; *ρ* is the density of water; *U*_∞_ is the uniform relative flow speed; *S*_w_ is the wing planform area (Table 1). Note that we chose the default parameters of liquid water in the flow solver (ANSYS Fluent), which are slightly different from the parameters of seawater. However, all the analyses across different wing posture (sweep angles, angles of attack) or relative flow speeds will be made in the normalised form (lift or drag coefficient); therefore, the slight variation in fluid parameters should not alter the conclusion. The forces were averaged over 1000 time steps after the force fluctuation diminished or became cyclic. The wing area was obtained by projecting the planform at feathering angle = 0° to the XY-plane.

#### Force balance estimation

To examine the obtained forces in the more realistic swimming situation, the vertical force balance was considered. For simplicity, the following force balance equation for the vertical direction is considered:

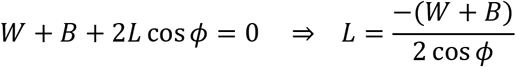

where *W* is the whole-body weight, *B* is the buoyancy, *L* is the lift from a single flipper, and *ϕ* is the anhedral angle (figure 2). The lift from the body is assumed to be negligible. In the real penguin, this would correspond to the situation where a penguin had flapped its wings then stopped for near-level gliding. Other situation such as ascending or descending glides are not considered. From this equation, we can obtain the lift coefficient of a single flipper which balances the vertical forces, *C*_L,equ,*ϕ*_ as:

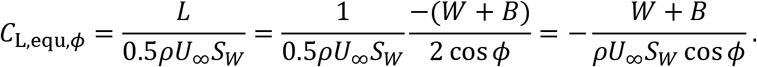

**Figure 2.**
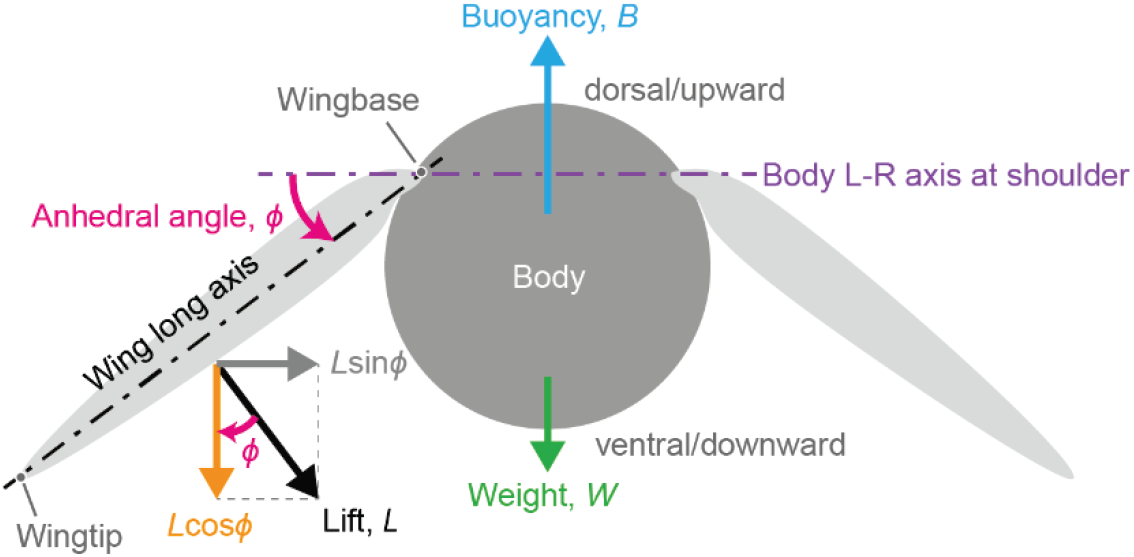
Frontal-view schematics of forces to explain the vertical force balance.

The mass of the penguin was measured with a scale on the same day before the swimming. The buoyant force was estimated based on a side-view photograph of the body for another individual and scaled with the flipper length assuming the isometric relationship between the flipper and body volume. The cross-sectional shape of the body was assumed to be circular. The three-dimensional body model generation and the volume calculation was done with Rhinoceros 5. The weight is defined as *W* = −*Mg* where *M* is mass and −*g* is the gravitational acceleration pointing downward. The body mass was 6.54 kg (same individual as the flipper 3D scan, though on a different day). Assuming *g* = 9.807 m s^−2^, *W* = −64.1 N where negative means downward. The buoyancy *B* was estimated to be 68.6 N, based on an approximate 3D model of the body, which was made from a side-view photograph of another individual assuming the circular body cross-section and scaled with the flipper length.

### Water tunnel test for validation

A water tunnel experiment was performed to validate the CFD results. A 1/2-scale wing model was created with CAD software (Fusion 360, Autodesk, Inc., USA) based on the same CAD model for the CFD and 3D printed with the layer thickness of 15 μm (AGILISTA-3100, Keyence, Japan). The base of the wing was attached to a 2-axis force sensor (LMC-21426-10N, Nissho Electric Works, Japan) to measure the lift and drag at various pitch (feathering) angles. The wing was inserted from a slit (60 mm × 30 mm) on a cover plate for the test section (1000 mm × 300 mm × 200 mm) of a circular-type water tunnel (Personal tank PT-100-mod, West Japan Fluid Engineering Laboratory Co., Ltd., Japan). The wing was held vertically from above so the wingtip points downward. This corresponds to the sweep angle *Λ* = 0°. The flow speed was set at 2.0 m s^−1^. The mean chord length was 27.5 mm and the water temperature was 17.1 °C. The corresponding Reynolds number was therefore 5.1×10^4^. The wing length from wingbase to wingtip was 127 mm but approximately 10% of the wing length near the wingbase was not submerged in the water. The wing area (planform area) that were submerged in the water was 3.27×10^3^ mm^2^. The sampling rate and the duration of the force measurement for each angle of attack was 10,000 Hz and 10 s, respectively. The average forces over 10 s were used to calculate the lift and drag coefficients. The lift coefficient and drag coefficient plots were shifted in the x-axis (angle of attack) direction so that the zero-angle of attack is defined as the zero-lift angle.

## Results

### Wing angles in gliding

In total, 78 horizontal forward gliding sequences were collected. In summary, the flap angle decreases with swimming speed (i.e., the anhedral or negative dihedral increases), while the sweep angle and angle of attack slightly increase with swimming speed (figure 3).

**Figure 3.**
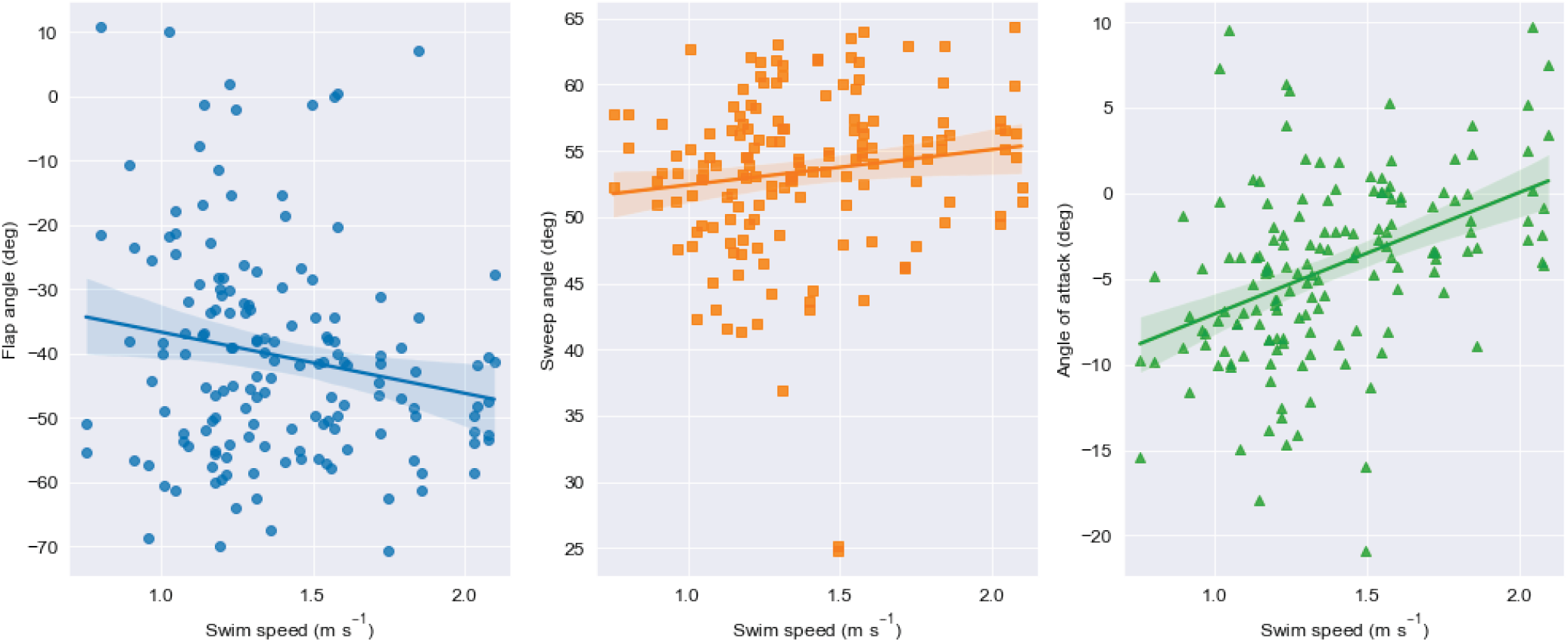
Wing angles against swimming speed in forward horizontal gliding. Flap angle (A), sweep angle (B), and angle of attack (C). Each symbol represents the angle of the left or right wing for each gliding bout. The lines and bands are linear regression and 95% confidence interval drawn with regplot command in seaborn(Waskom, 2021).

### Validation

The experimental and numerical results show good agreement in lift coefficient *C*_L_ (figure 4A). There is a slight deviation in *C*_L_ at high negative angles of attack. The drag coefficients *C*_D_ also show the overall good agreement, but beyond the stall angles (at around −15° and +15°) the *C*_D_ in the experiment is greater. In the experiment, the approximately 10% wing length from the wingbase was assumed not to have been immersed in the water, but the slight wave might have increased the actual surface area that was affected by the water and resulted in the slightly greater *C*_L_ and *C*_D_.

**Figure 4.**
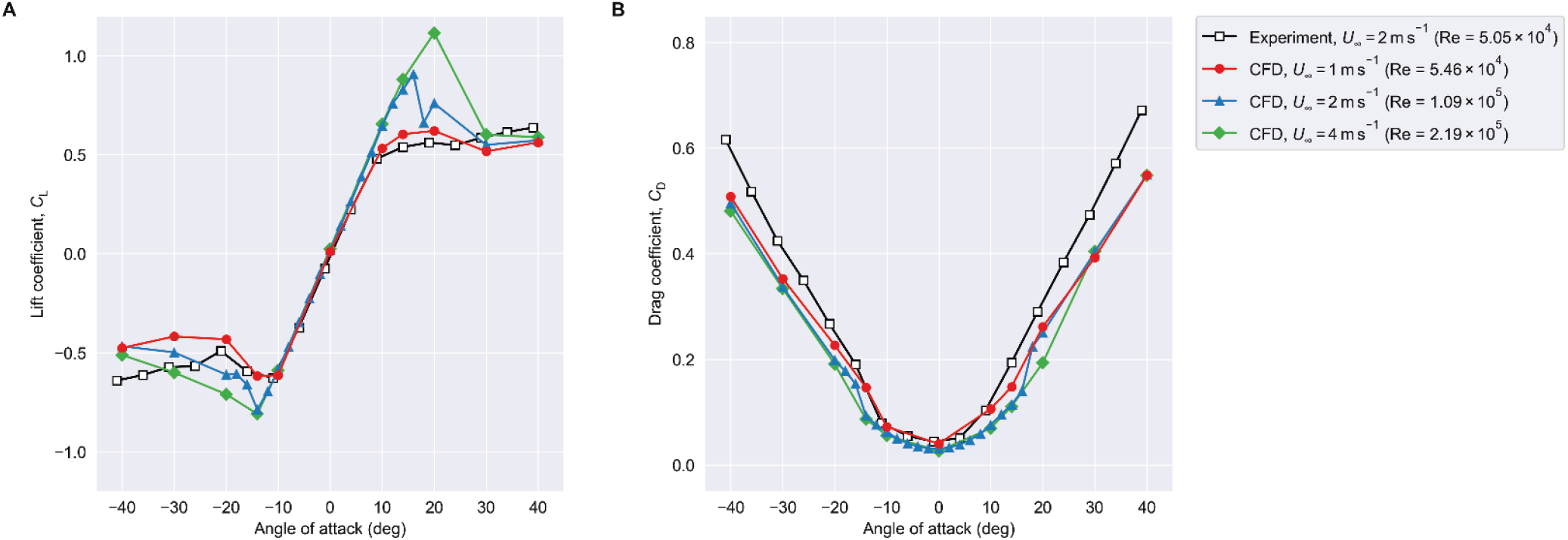
Comparison of lift and drag coefficients for the sweep angle *Λ* = 0°. Keys: open black squares, experiment with *U*_∞_ = 2 m s^−1^ (Re = 5.05×10^4^; note the 3D printed wing model is half-scale of the real wing); red circles, CFD with *U*_∞_ = 1 m s^−1^ (Re = 5.46×10^4^); blue triangles, CFD with *U*_∞_ = 2 m s^−1^ (Re = 1.09×10^5^); and green diamonds, CFD with *U*_∞_ = 4 m s^−1^ (Re = 2.19×10^5^). Sweep angle *Λ* = 0° for all the cases.

### Forces

The lift slope around *α* = 0° is greatest in the no-sweep wing (figure 5A, blue circles). Increasing the sweep angle to 30° slightly reduces the slope (figure 5A, orange triangles) but the change is small. In contrast, sweeping by 60° substantially reduced the slope (figure 5A, green squares). The stalling tendency is also dependent on the sweep angle. The wing with 0° sweep shows the clear stall beyond −14° and +16°, and the magnitude of lift coefficients beyond these angles rapidly decreases towards α = ±40°. In contrast, when the sweep is 30°, the stall occurred much gradually, in particular for the negative angles of attack. When the sweep is 60°, the change in force is even more gradual and there is no clear stall angle.

**Figure 5.**
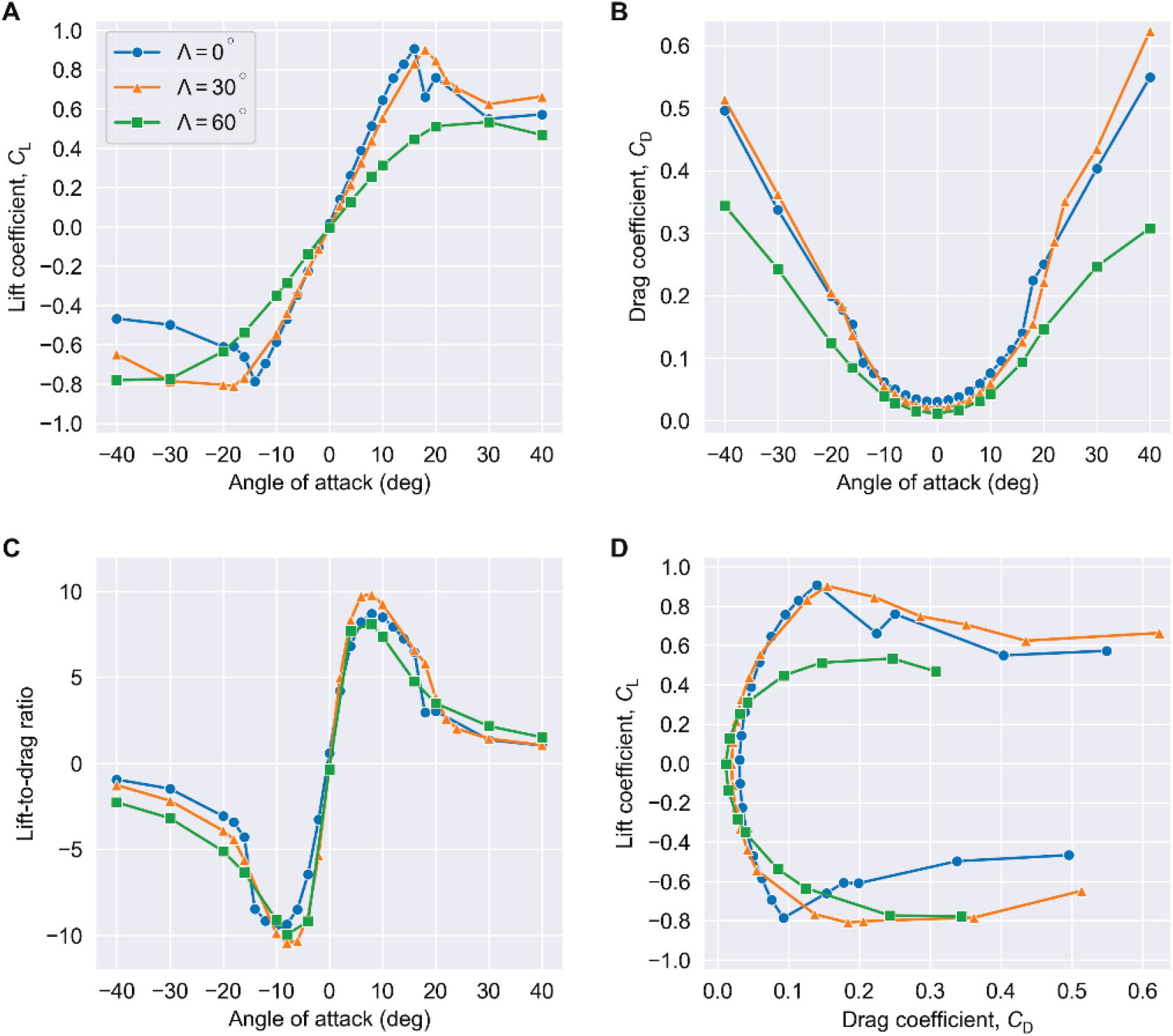
Hydrodynamic forces of a penguin flipper gliding at *U*_∞_ = 2 m s^−1^. The variation of lift coefficient (A), drag coefficient (B), and lift-to-drag ratio (C) against the change in angle of attack *α*. The drag polar (D). Keys: blue circles, sweep angle *Λ* = 0°; orange triangles, *Λ* = 30°; and green squares, *Λ* = 60°.

The drag coefficients for the sweep angles of 0° and 30° are overall quite similar, while when the sweep is 60°, substantially smaller drag coefficients across the range of angles of attack can be seen, in particular the pre-stall angles of attack. For 0° and 30° sweep cases, a discontinuous increase in drag coefficients can be seen at around stall angle. Similarly to lift, no such abrupt change can be observed for the drag at 60° sweep. The side-force coefficient shows the inverse tendency (figure 5B) compared to the drag coefficients, where the 0° sweep wing has nearly no side force but as the sweep increases the force coefficients increases for the higher angles of attack.

The maximum lift-to-drag ratio (*L*/*D*) magnitude of approximately 10 is achieved for 30° sweep wing at *α* = ±6– 8°, which are slightly before the stall (figure 5C). The overall values and the trends are similar across the range of sweep angels. It is interesting to see the similar trend of *L/D* can be seen even when the sweep is 60°, despite the marked deviation in the lift and drag coefficient from the lower sweep angles. In fact, at the higher angles of attack (*α* < −20° and *α* > 20°), the best *L*/*D* is achieved when the sweep the highest (*Λ* = 60°; green squares in figure 5C).

The drag polars (figure 5D) may be the most interesting plots of all the data we obtained. They indicate that for the pre-stall angles of attack, the sweep angle that can achieve the highest *L/D* is dependent upon the angle of attack. When the angle of attack magnitude is small, choosing the high sweep angle (*Λ* = 60°, green squares in figure 5D) can reduce the drag coefficient while keeping the (although small) lift coefficient. At an intermediate angle of attack magnitudes, one can attain higher *L/D* by reducing the sweep (*Λ* = 30°, orange triangles in figure 5D). At an even higher (but pre-stall) angle of attack magnitudes, no-sweep is the best (*Λ* = 0°, blue circles in figure 5D). This will be more clearly seen in the enlarged view near *α* = 0° (figure 6B).

**Figure 6.**
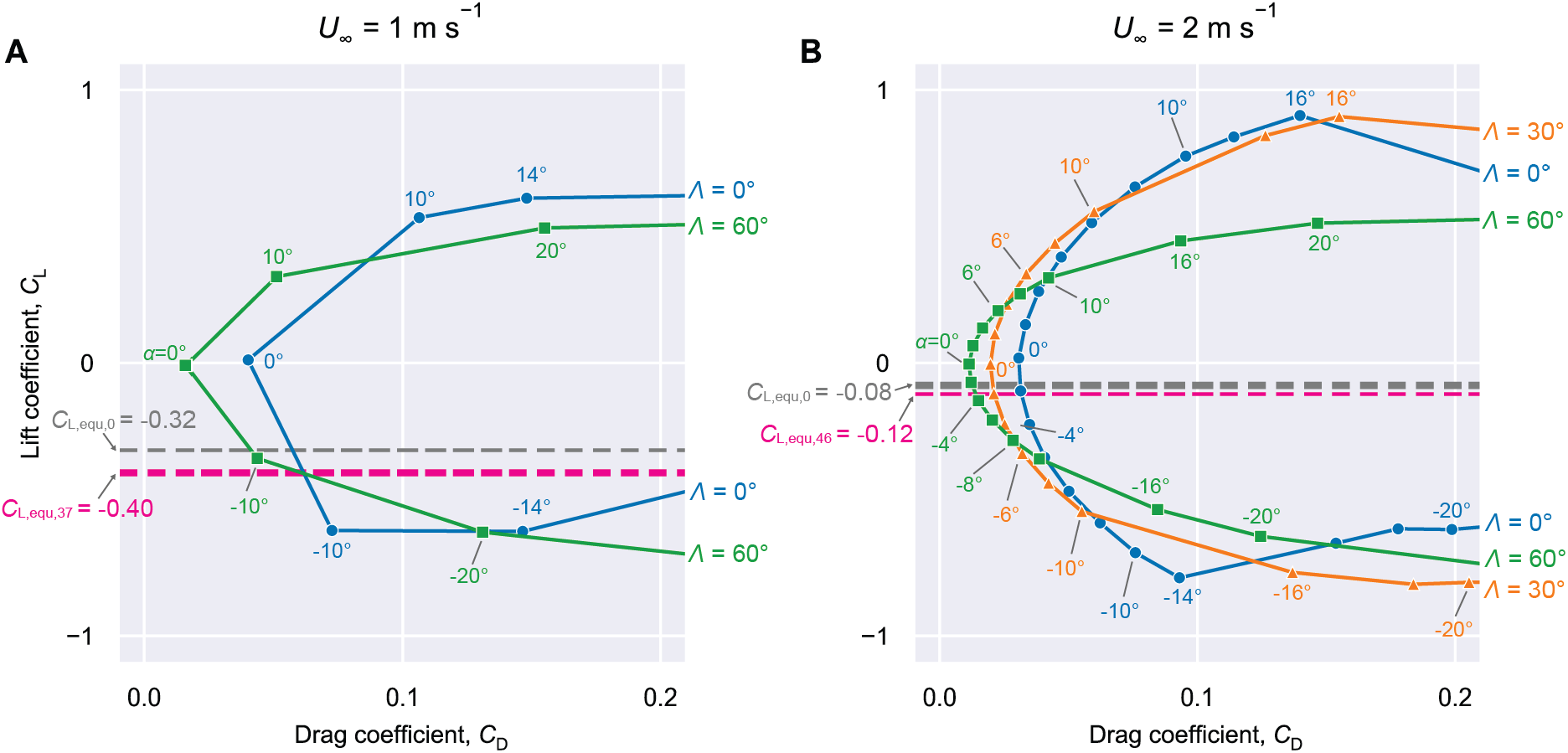
Enlarged view (−1 ≤ *C*_L_ ≤ 1, *C*_D_ ≤ 0.2) of the drag polars indicates the possible selection of the sweep angle according to the swimming speed. Swimming speed *U*_∞_ = 1 m s^−1^ (A) and *U*_∞_ = 2 m s^−1^ (B). Data series correspond to different sweep angles (blue circles, *Λ* = 0°; orange triangles, 30° (only for *U*_∞_ = 2 m s^−1^); and green squares, 60°). Grey and pink dashed lines correspond to the lift coefficients required for vertical equilibrium, where grey is assuming zero anhedral while pink is assuming realistic anhedral (shown as subscripts to the lift coefficients in degrees) from the regression.

### Force balance

To assess how the penguins should choose the sweep angles during gliding, we plotted an enlarged view of the drag polars for two swimming speeds: 1 and 2 m s^−1^ (figure 6), where only the region that is close to the zero-angle of attack was focused. In each plot, the lift coefficient that a single flipper must produce to balance the vertical forces is also plotted with two assumptions on the anhedral angle: zero anhedral (*C*_L,equ,0_, grey dashed lines) and realistic anhedral from the regression (*C*_L,equ,*ϕ*_, pink dashed lines). From the regression, the expected wing angles at the swimming speeds of 1 and 2 m s^−1^, respectively, are: anhedral angle (negative flap angle) *ϕ* = 36.7° and 46.3°; sweep angle *Λ* = 52.4° and 55.1°; angle of attack *α* = −7.05° and 0.03°. As expected, the anhedral slightly increases the required lift coefficient.

## Discussion

### Drag can be reduced by adjusting sweep angle with swimming speed

A gliding penguin would inevitably lose momentum due to the drag, and the swimming speed would consequently decrease. This will need to be compensated by the subsequent flaps to maintain the average swimming speed over the flap-glide cycles. It is therefore ideal to minimise the drag in gliding while generating sufficient vertically downward force component to counteract the positive buoyancy. We can obtain this optimum by examining the drag polar plot (figure 6). The horizontal dashed lines are the required lift coefficients, so we should simply find the leftmost (least drag) point that the horizontal line and the drag polar curve cross. At the swimming speed of 2 m s^−1^ (figure 6B), the magnitude of the lift coefficient required is small, and so is the angle of attack magnitude. Therefore, choosing a high sweep angle (*Λ* = 60°) is the best among the three sweep angles. If, for some reason, a much greater downward lift coefficient is required (i.e., horizontal line shifts downward in the plot), then it is possible that reducing the sweep angle and increasing the angle of attack magnitude would result in better glide performance. This can happen when the body weight decreases or buoyancy increases, or both. On the other hand, at a lower swimming speed of 1 m s^−1^ (figure 6A), the high sweep angle does not seem to be the best option. The pink dashed line crosses both the high and low sweep angle curves, indicating the intermediate sweep angle (0° < *Λ* < 60°) would be the best. In fact, the sweep angles and angles of attack predicted from the regression for both swimming speeds are showing the same trends (figure 3B,C) and the values are not far off, which supports that our analysis is sound. It should be noted that a similar adjustment of sweepback has been found in a gliding bird, swift (figure 2d in Lentink et al., 2007), where the authors described this phenomenon “enlarges the aerodynamic performance envelope of swift wings.” We suggest the same can be said to the penguin flippers while gliding.

Interestingly, from figure 6 we can see that increasing the amount of anhedral (negative dihedral, or negative flap angle) increases drag: i.e., the point that the pink dashed line (nonzero anhedral wing) crosses with the drag polar curve is always right of the point that the grey dashed line (zero-anhedral wing) crosses the same polar curve (figure 6). Why do the penguins choose such hydrodynamically less efficient wing posture of nonzero anhedral then? One possibility is roll stability. In aeroplanes, it is well known the dihedral (wingtips being above the wingbases) gives the roll stability, which is called the dihedral effect. In penguins, however, the buoyancy is generally positive, and the vector sum of the buoyancy and weight is pointing upward. The vector is essentially equivalent to the gravitational acceleration vector in the aeroplane but acting in the opposite direction. Therefore, in penguin, the wings with dihedral would result in roll instability and wings with anhedral should result in roll stability. The moderate anhedral angles we observed in the penguins may therefore be contributing to the roll stability *via* this “anhedral effect.” The increasing trend of the anhedral (negative flap angle) with swimming speed (figure 3A) may support this as well, as the higher the speed the stronger roll stability may be preferred. This anhedral effect is expected to be universally used by the other wing-propelled swimmers that have positive buoyancy (e.g., mammals, birds, or turtles), and it seems that many of them are indeed having some anhedral angle while gliding.

Pitching torque balance is another factor that was not explored in the current study but would be important in choosing the gliding wing posture. If the sweep angle is very small or even negative (i.e., forward-swept), the centres of pressure of the wings locate anterior to the centre of mass, which would generate the nose-down pitching torque because of the downward force component. Increasing the sweep angle would reduce the moment arm (distance between the centre of pressure and the centre of mass) and decrease the pitching torque.

### Flow visualisation

Flow visualization by surface pressure and limit streamlines (figure 7) indicates that the flow is generally attached on the dorsal side at negative angles of attack, but detached near the leading edge. With a low sweep angle, the leading-edge vortex (LEV, or laminar separation bubble, LSB) seems to have burst at a high (post-stall) angle of attack (figure 7B). In contrast, with a larger sweep angle, the leading-edge vortex is attached even at a high angle of attack (figure 7D), and likely to prevent an abrupt stall. It is also noted that with a high sweep angle, the flow on the wing’s proximal region is largely attached, which is likely to contribute to the substantially lower drag.

**Figure 7.**
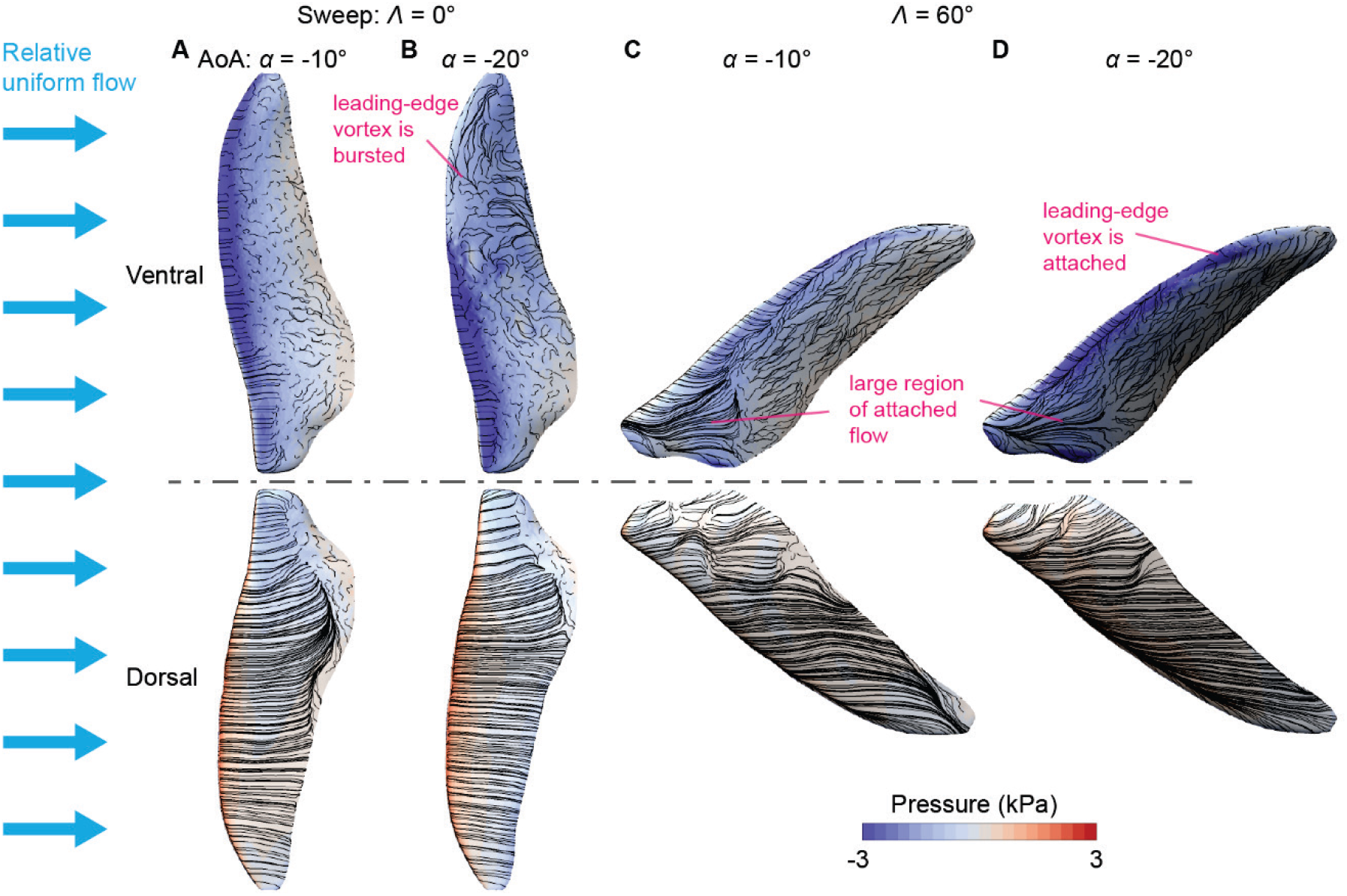
Surface pressure and limit streamlines show the contrasting flows across glide angles. Sweep angle *Λ* = 0° (A, B) and 60° (C,D). Angle of attack *α* = −10° (A, C) and −20° (B, D). The relative flow is from left to right.

### Limitations and future directions

The actual flippers are not rigid, but flexible structures. The dynamic wing deformation is evident in flapping (Harada et al., *under review*) but even in gliding the wing may be deformed and differ from the wing shape in the air. It is also possible that the wing deforms adaptively for each angle of attack. Such wing deformation needs to be explored in future studies. It would also be an interesting direction to explore whether the micro-roughness emerging from the filament-like feathers has a similar benefit as in the sharkskin (riblet in general).

We are aware that the body may have a non-negligible effect on the wing performance. In particular, when the sweep angle is high and the proximal trailing edge is close to the body surface, physical or hydrodynamic interfere are likely to occur. By physical interference we mean the feathering angle of the wing may be restricted to a certain range at the high sweep, and by hydrodynamic interference, we mean the flow is affected by the proximity of the body surface. Treatment of such complexities is beyond the scope of the current study, but it would be worth pursuing in future study.

It has recently been proposed to design or modify the shape and attachment location of the logging devices so they will have a less hydrodynamic negative effect (less increase in drag, etc.) on animals (e.g., Fiore et al., 2017; Kay et al., 2019). A similar approach may be employed for improving the flipper band design, which is generally considered to have a negative impact on penguins (e.g., Saraux et al., 2011).

## Conclusions

We performed the computational fluid dynamics study on a penguin flipper under gliding condition. It was found that the wing sweepback is effective in preventing the abrupt wing stall. The maximum lift-to-drag ratio was achieved at the intermediate sweepback. Most notably, the drag polars across the range of sweepback angles suggested the expanded swimming envelope and the potential importance in the selection of the optimal sweepback depending on the swimming speed.

## Acknowledgements

This work was supported by the Grant-in-aid number JP18H05468. We thank Nagasaki Penguin Aquarium for their support in the experiments and measurements using penguins. We thank Yusuke Iwasaki for his help in scanning the penguin flipper. We thank Hiroki Kayasuga for his help with the water tunnel experiment. The numerical simulation was carried out using the TSUBAME 3.0 supercomputer at Tokyo Institute of Technology.

